# Basal Ganglia Stimulation Ameliorates Schizophrenia Exploration Anomalies

**DOI:** 10.1101/2023.08.13.553111

**Authors:** Nir Asch, Noa Rahamim, Uri Werner-Reiss, Zvi Israel, Hagai Bergman

## Abstract

Any learning agent must balance between exploiting its knowledge and exploring new alternatives. Schizophrenia patients are known to have maladaptive exploration-exploitation (E-E) balance^1,2^ and are impaired at reversal learning tasks as early as their first psychotic episode. The cortico-basal ganglia (BG)-dorsolateral prefrontal cortex (DLPFC) network plays a significant role in learning processes^3,4^. However, how this network maintains the E-E balance and what alters the balance in schizophrenia remains elusive. Using a combination of extracellular recordings, pharmacological manipulations, macro-stimulation techniques, and an adaptive reinforcement learning model, we show that in the non-human primate (NHP), the external segment of the globus pallidus (GPe, the central nucleus of the BG network) maintain this balance. Furthermore, whereas the chronic, low-dose administration of N-methyl-D-aspartate (NMDA) receptor (NMDA-R) antagonist, phencyclidine (PCP) leads to E-E imbalance, low-frequency GPe macro-stimulation restores it. E-E balance provides a holistic framework to resolve some of the apparent paradoxes that have emerged within schizophrenia research^2^. Our findings suggest that Schizophrenia symptoms may reflect abnormal DLPFC-BG E-E balance, and GPe stimulation may be advantageous for these patients.

## Main

Humans and NHPs are highly developed learners, a trait with great evolutionary benefit. One of the challenges for optimizing learning is navigating one’s behavior between exploiting its knowledge and exploring new, possibly better, alternatives. Exploration is further divided into two categories: Directed exploration, driven externally by a mismatch between predictions and outcomes, allowing the agent to modify its behavior in light of experience, and random exploration, internally driven without such mismatch^5–8^. Random exploration is crucial in optimizing behavior, particularly when encountering local maxima, where the agent may possess an inferior solution to a problem while a better solution exists.

The DLPFC and the BG networks are well integrated^9–11^. They are recognized as the major underlying neural circuitry of goal-directed (governed by the cortico-cortical network) and implicit/habitual (controlled by the basal ganglia network) learning paradigms^3,12–14^. Nonetheless, the neural underpinnings keeping the E-E balance still need to be elucidated. The GPe (the central nucleus of the main axis of the BG network, connected to both input and output BG layers)^14–16^ and the DLPFC^7,17,18^ have been hypothesized as possible hubs for controlling the E-E balance. The BG has also been suggested as having an active gating mechanism dynamically regulating the influence of incoming stimuli on the cortical working memory (WM) system^19–21^.

Schizophrenia patients exhibit impairment in reversal learning tasks (directed exploration, corresponding with schizophrenic negative and cognitive clinical symptoms)^22^, and increased random exploration (corresponding with the positive symptoms)^1^. They have impaired WM associated with DLPFC dysfunction^23,24^ and a hyperactive and enlarged GPe in correlation with the severity of their symptoms^25,26^. Therefore, their increased tendency for random exploration may result from an over-active ‘exploration generator’ (e.g., the GPe)^27^ or their WM deficit. To gain deeper insights into the neural mechanisms underlying E-E balance in both healthy and schizophrenia-affected states, we employed a chronic administration of the NMDA-R antagonist PCP to induce a schizophrenia-like state^28–32^ in two African green monkeys (*Chlorocebus aethiops sabaeus* (vervet), females, weights: ∼4 kg). These monkeys were extensively trained in a reversal-learning task, and their GPe and DLPFC activities were recorded during task performance (Fig. 1a-b). By examining the neural dynamics of the DLPFC and GPe in both the naïve and PCP-induced schizophrenia model, we aimed to investigate the neural basis of E-E control.

**Figure 1:**
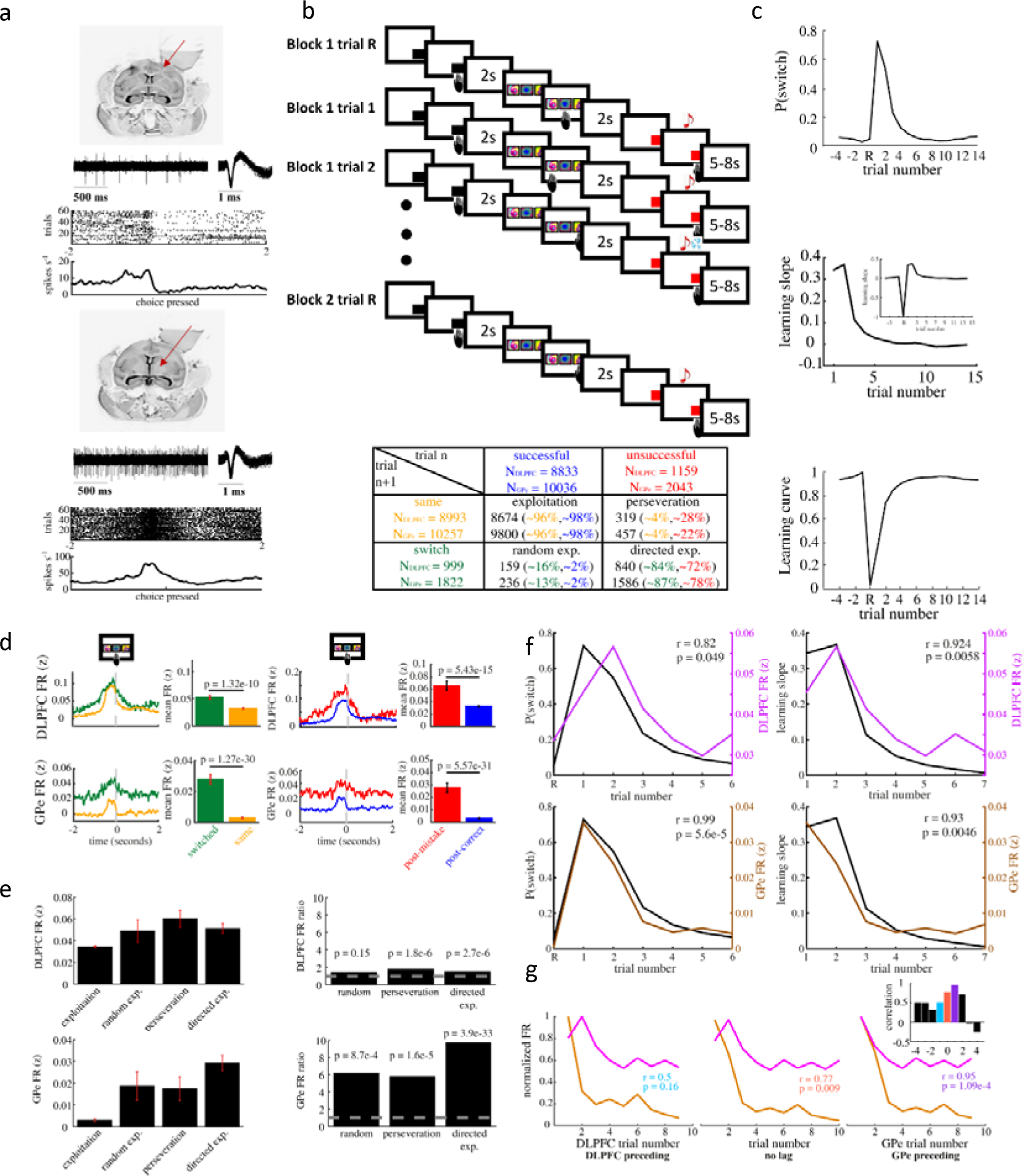
GPe activity correlates with exploratory actions, while DLPFC activity is associated with the learning slope and lags behind GPe activity changes. **(a) Top** - MRI of the NHPs’ brain and recording chamber. Red arrow points to the DLPFC. Middle, extracellular recordings of DLPFC exemplary neuron (left) and 100 randomly chosen superimposed spike waveforms of the recorded cell (right). Bottom, Raster display and a PSTH of the neuron’s firing during 60 trials around choice selection. **Bottom** – The same as the top image, but for the GPe. **(b) Top** - Reversal learning task design. Monkeys had to identify hidden rule changes and learn the new rule. The first line represents a reversal trial (R) in which the monkey is unaware of the change of the new rule and therefore chooses the wrong cue. The second line represents the next trial in the block in which the monkey changes its choice to another incorrect choice, and in the third line, the monkey changes again, this time receiving a reward. The fourth line represents the following block in which the rule is changed without the NHPs’ knowledge. The NHP then chooses the same stimulus as before, but this time receives no reward. **Bottom** – A table showing the amount (and fraction) of all neural recorded trial types of the two brain regions. Text color coding indicates the fraction out of recorded cells within each category from the total sum. For example, our neural data set of perseveration trials consists of 319 DLPFC and 457 GPe recordings, constituting 28% and 22% of all neurally-recorded unsuccessful trials, respectively, and 4% of all the neurally-recorded same-choice trials. **(c) Top** – Switch probability. **Middle** – The learning slope (i.e., the derivative of the learning curve). Inset – the learning slope across blocks. **Bottom** – The learning curve. ‘R’ indicates the reversal trial, trials prior/post to reversal are of negative/positive sign, respectively. **(d) Left** – DLPFC and GPe firing rate around choice selection, comparing switched trials (green) and not switched trials (orange) using the two-sample t-test. **Right** – DLPFC and GPe firing rate around choice selection, comparing exploration-favorable trials (post-mistake, red) and exploitation-favorable trials (post-correct, blue) using the two-sample t-test. **(e) Left** – mean and SEM DLPFC and GPe firing rate leading to stimulus choice selection of the four trial types. **Right** – firing rate ratio between the firing rate leading to stimulus choice selection of the three non-exploitation trial types with the firing rate leading to choice selection in exploitation trials. Ps represent the p-values corresponding to the two-sample t-test between the FR of each of the three behavioral types (random exploration, perseveration, and directed exploration) and exploitation (p-values are Bonferroni corrected for multiple comparisons) **(f) Left** - Mean DLPFC (top) and GPe (bottom) firing rate of the two seconds preceding choice selection (purple) with the NHPs’ probability to switch to a new key (black line). Correlation values and their corresponding p-values between switch probability and neural activity are represented by ‘r’ and ‘p’ respectively. **Right** - Mean DLPFC (top) and GPe (bottom) firing rate of the two seconds preceding choice selection (purple) with the NHPs’ learning slope (black line). **(g)** Correlation between mean DLPFC and GPe firing rates (recorded during the two seconds leading to choice selection) during the first ten trials. **Left** - Comparing DLPFC activity in trial N with GPe activity in trial N+1. **Middle** – Comparing DLPFC and GPe concurrent activities. **Right** - Comparing DLPFC activity in trial N+1 with GPe activity in trial N. Inset - correlation values between DLPFC and GPe activities calculated from -4 (DLPFC activity precedes GPe activity by four trials) to +4 (GPe activity precedes DLPFC activity by four trials) trials lag. **See also extended data figures 1-4**

### Naïve behavior and associated neural activity patterns

The NHPs were trained to acquire and reverse their response to novel visual cues in a between-block design (Fig. 1b). First, they had to identify the rule associated with reward outcome. Once learned, the rule changed after reaching a predefined success criterion (cutoff selected randomly for each block to avoid identification of the rule), and a new block began without any external cue (average of 18.8 trials [ranging between 13-40] per block, a total of 1,673 blocks, for the naïve state). Thus, in the first trial of every block (reversal, R, trial), the NHPs experienced a prediction-outcome mismatch in which choosing the previously correct cue elicited no reward. Ideally, the NHPs should start directed exploration at the R+1 trial (Fig 1b). The NHPs showed fast learning dynamics, rapidly identifying the rule change, and initiating productive directed exploration (Fig. 1c). During directed exploration, the NHPs expressed well-functioning WM, avoiding previously selected erroneous cues in search of the new rule (Fig. 1c and extended data Fig. 1a,b). An optimal learning agent would find the correct response with an average of 1.5 trials. The NHPs found the cue associated with positive reward outcome within 3.11 ± 0.04 trials (mean ± SEM). Once identifying the new correct cue, the NHPs were able to exploit their newly acquired knowledge to maximize their gain, completing the learning phase (defined as three sequential successful trials) within 5.44 ± 0.05 trials. Post-learning, the NHPs achieved a mean success rate of 93% during the learning plateau (Fig. 1c). Their response times further reflected their cognitive effort, lengthening during the learning phase with directed exploration trials and decreasing thereafter (extended data Fig. 1a).

To better understand the underlying neural mechanism of the E-E balance, we analyzed the neural dynamics around choice selection in the DLPFC and GPe. Assessing the neural activity leading to exploratory (i.e., switch of choice) and non-exploratory (same choice) actions (Fig. 1d, and extended data Fig. 2), we found that the firing rate in both regions increased before exploratory compared to non-exploratory decisions. To assess the neural activity leading to directed vs. random exploration, we compared the average neural discharge rate leading to stimulus choice selection in post-mistake and post-correct trials. We found that both regions elevated their firing rate in post-mistake compared to post-correct trials (Fig. 1d). Like all biological creatures, our NHPs were imperfect. They did not explore after every prediction-outcome mismatch (a switch occurred after 78.2% of mistakes) or exploit after every congruent prediction outcome (a switch occurred after 2.25% of corrects). We, consequently, analyzed the neural activity leading to a switch and non-switch in post-mistake trials (directed exploration and perseveration, respectively, Fig. 1e and extended data Fig. 2b). The average firing rates before choice selection in directed exploration and perseveration trials were not significantly different in either brain region. Similarly, we analyzed the neural activity leading to a switch and non-switch in post-correct trials (random exploration and exploitation, respectively, Fig. 1e and extended data Fig. 2c). DLPFC activity showed the same pattern with no significant difference between random exploration and exploitation. Conversely, the GPe showed a significantly higher firing rate before random exploration than exploitation trials.

Whereas both regions increased their activity after a prediction-outcome mismatch (Fig. 1d), neither convincingly modified its activity in directed exploration compared to perseveration. We, therefore, conducted a more detailed analysis comparing the neural dynamics to the behavioral ones during the learning phase (Fig. 1f and extended data Fig. 3). We found GPe activity to be highly correlated with switch probability, while DLPFC activity correlated only marginally. Switch probability (especially in the early part of the block) is strongly associated with the prediction-outcome relationship of the previous trial. Nonetheless, while directed exploration is driven by the prediction-outcome mismatch of previous events, its direction is determined by new predictions. Thus, learning speed is determined by these two parameters, initiation of exploration (i.e., switch probability) and predictions. The faster directed exploration is initiated, and the more accurate the predictions are, the faster learning becomes. We, therefore, compared the neural activity with the learning slope (i.e., the derivative of the learning curve), representing learning speed. Whereas both regions neural activity correlated similarly with the learning slope, the DLPFC exhibited a much better correlation with the learning slope than with switch probability (Fig. 1f).

A switch can either be successful or unsuccessful. While choosing successfully in the first trial after reversal is sheer chance, choosing successfully later in the block demands higher WM effort, remembering previously chosen incorrect trials and predicting the location of the correct stimulus cue. Hence, we compared the correlation between both regions’ neural activity and all switch probability types: general, successful (associated with WM load), and unsuccessful switch probabilities (extended data Fig. 3b). The DLPFC was highly correlated with the likelihood of making a successful switch and did not correlate with the chance of making an unsuccessful switch. The GPe, on the other hand, exhibited significant correlations with both successful and unsuccessful switch probabilities. These results suggest that DLPFC activity is associated with increasing memory load, whereas GPe activity is associated with exploratory behavior. Since exploratory behavior is strongly driven by previous reward outcomes, we tested whether reward outcomes are encoded in GPe activity and whether this information is carried across trials. Indeed, comparing GPe activity change induced by reward acquisition, we found that the GPe elevates its discharge rate in response to prediction-outcome mismatch and that this elevation persists in the subsequent trial (extended data Fig. 3c-d). This activity pattern was not observed in DLPFC activity (extended data Fig. 3c-d).

We, therefore, compared the two regions’ activity dynamics throughout the task. While their concurrent activities only marginally correlated (Fig. 1g), comparing their activities in subsequent trials (i.e., comparing the correlation between GPe activity in trial ‘n’ with DLPFC activity in trial ‘n+1’) exhibited a strong correlation (Fig. 1g). Furthermore, whereas GPe units responded similarly to the three choice stimuli (some cells increased while others decreased their activity), DLPFC units expressed a more sophisticated activity pattern, responding differently to each stimulus (extended data Fig. 4). In summary, these results show that the GPe increases its activity after prediction-outcome mismatch and with both directed and random exploratory actions. The DLPFC, on the other hand, increases its activity in subsequent trials associated with WM load and the prediction of the next optimal action process.

### PCP-induced behavioral and neural activity changes

To investigate the neural correlates of schizophrenia, we examined the impact of chronic, low-dose PCP treatment on neural activity and concurrent task performance. Analysis of PCP administration effects on spontaneous, background neural activity revealed baseline changes in both regions. DLPFC LFP beta activity and background firing rates were reduced under PCP administration. GPe activity, on the other hand, showed a mild reduction in beta activity but an increase in its background firing rate. GPe gamma activity showed a significant rise in a narrow bandwidth around 30-40 Hz, while DLPFC gamma activity remained similar to the naïve state. Interestingly, both regions’ gamma activity and background firing rates rose significantly post-PCP, suggesting a long-term and/or rebound effect (extended data Fig. 5 and 6). The coefficient of variation of the inter-spike intervals (a proxy for burst/periodic discharge pattern) remained relatively unchanged under all conditions in both regions. Additionally, the overall appearance of ‘pausing’ activity in GPe neurons continuously decreased from the naïve to chronic PCP to the post-PCP state (extended data Fig. 6).

Importantly, PCP had a detrimental effect on the NHPs’ ability to perform the task, resulting in slower learning and achieving a lower success rate plateau (Fig. 2a-d). The ability to successfully switch during the learning phase decreased to chance levels, not improving with task progression (Fig. 2b, inset). Consequently, the identification of the new correct response (extended data Fig. 7) and the achievement of the learning criterion (Fig. 2c) were delayed. Overall, the NHPs showed a slight decrease in directed exploration behavior and a significant increase in random exploration behavior (Fig. 2d). This contributed to a decrease in the learning plateau success rate. Furthermore, the response times for stimulus choice selection increased throughout the block, probably indicating the NHPs’ growing uncertainty or reduced motivation. Response times for claiming reward outcomes no longer correlated with task performance (extended data Fig. 7d,e). Finally, following the cessation of PCP treatment (i.e., recording conducted at least two weeks later), the NHPs’ performance improved across all parameters (Fig. 2 and extended data Fig. 7).

**Figure 2:**
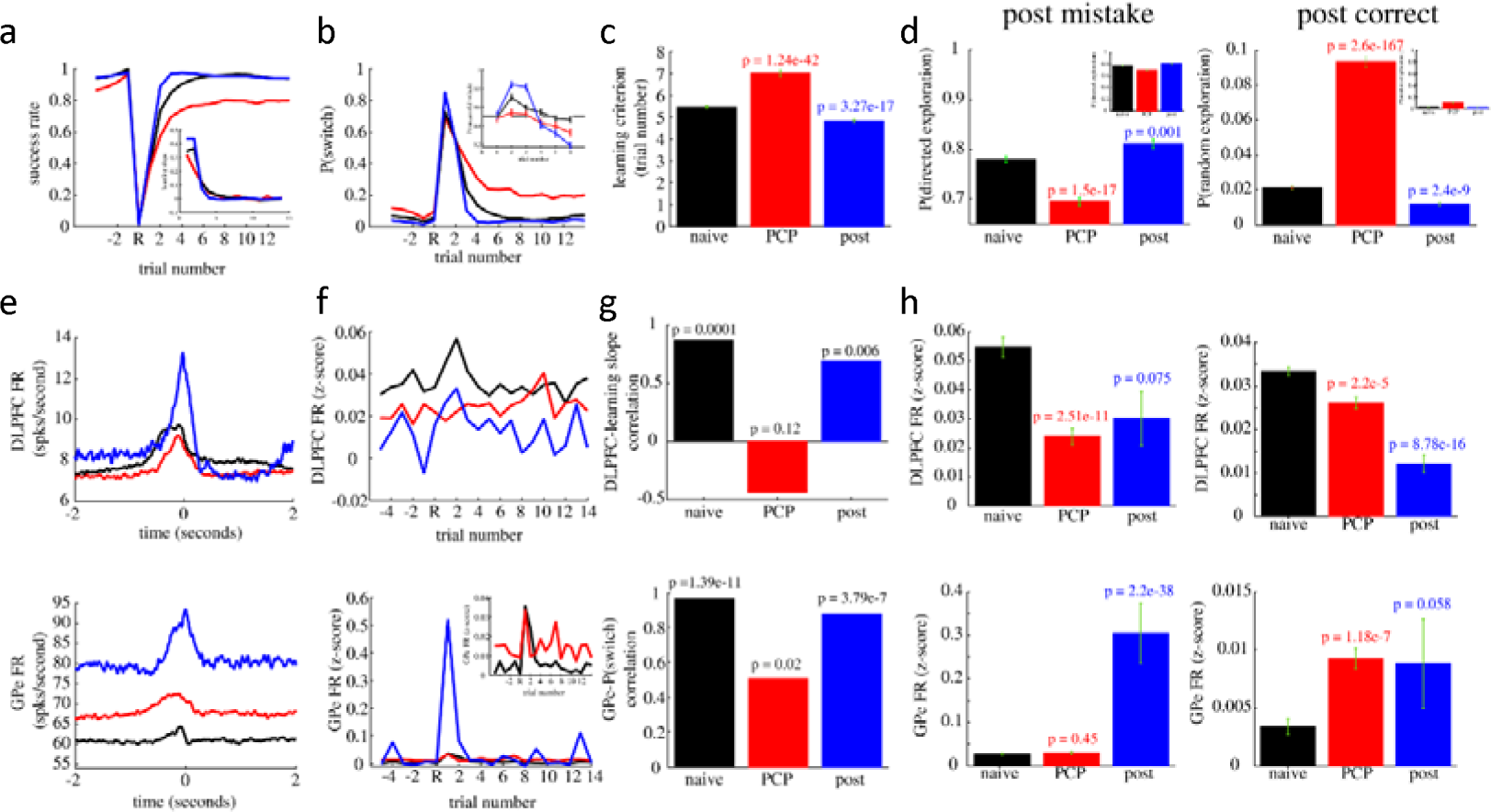
PCP induces decoupling of DLPFC activity from learning dynamics, increases GPe activity, and impairs exploration patterns. **(a)** The NHPs’ learning curve under all conditions naïve (black), under PCP administration (red), and post-PCP (blue). Inset shows the associated learning slopes. **(b)** Average switch probability throughout the task. Inset shows the NHPs’ probability of choosing the correct stimulus choice given that a switch was made during the first six trials (chance level is marked with a dashed grey line). **(c)** Learning criterion, the average, and SEM of the trial number in which learning was achieved (learning defined after achieving three consecutive successful trials). Ps represents the p-value of the two-sampled t-test comparing PCP and post-PCP values to the naïve result. P-values are Bonferroni corrected (i.e., p-values are multiplied by the number of multiple comparisons). This annotation follows in all subsequent plots. **(d) Left -** Directed exploration probability (i.e., switch probability post mistake - the probability that the NHPs switched their choice after an unsuccessful trial). **Right –** random exploration probability (i.e., switch probability post correct - the probability that the NHPs’ switched their choice after a successful trial). **(e-h)** analysis of neural activity throughout the task (the number of naïve GPe and DLPFC recorded trials were 12,079 and 9,992, respectively. The number of PCP GPe and DLPFC recorded trials were 8,291 and 5,034, respectively. The number of GPe and DLPFC post-PCP recorded trials were 1,788 and 1,858, respectively). **(e)** DLPFC (top) and GPe (bottom) average firing rate around choice selection (at time zero). **(f)** DLPFC (top) and GPe (bottom) mean firing rate (z-scored) during the two seconds leading to stimulus choice selection throughout the task in the naïve state (black), under PCP administration (red) and post-PCP (blue). **(g) Top -** Correlation between DLPFC activity and the learning slope (across the entire task). **Bottom –** correlation between GPe activity and switch probability (across the entire task). **(h) Left -** Mean and SEM of DLPFC (top) and GPe (bottom) activity leading to stimulus choice selection in trials following an unsuccessful trial (i.e., post-mistake) comparing naïve results (black) to the PCP (red) and post-PCP (blue) conditions. **Right** – the same as the left, comparing trials following a successful trial (i.e., post-correct). **See also extended data figures 5-8**

Comparing the neural and learning dynamics throughout the task allowed for a complementary understanding of the PCP-induced neuro-behavioral changes. DLPFC activity leading to stimulus choice selection slightly decreased in the chronic PCP stage and increased post-PCP (Fig. 2e). GPe activity around stimulus choice selection increased with PCP administration and further increased post-PCP (Fig. 2e). The DLPFC activity under PCP did not correlate with the learning slope, but this correlation was regained post-PCP (Fig. 2f,g). At the same time, GPe activity remained correlated with the probability of making exploratory actions under the influence of PCP, albeit with a slightly reduced correlation value, regaining its high-value post-PCP (Fig. 2f,g).

Next, we compared the activities of both regions leading to stimulus choice selection in post-mistake and post-correct trials (Fig. 2h). We found that DLPFC activity in both trial types decreased and, notably, the neural activity was no longer discriminative between the two (Fig. 2h, and extended data Fig. 7f). Conversely, GPe activity remained discriminative between the two trial types (Fig. 2h and extended data Fig. 7f). While GPe activity during post-mistake trials (i.e., after prediction outcome mismatch, directed exploration driving force) showed a non-significant increase, GPe activity in post-correct trials increased fourfold corresponding with the increase in random exploration probability. These neuro-behavioral changes in GPe activity and exploration patterns may explain the reduced correlation between switch probability and GPe activity in the schizophrenia PCP-modeled state.

Discontinuation of PCP administration restored the NHPs’ learning and neural dynamics towards those observed during the naïve state (Fig. 2). Concomitantly, gamma oscillatory activity, known to be associated with attention and WM, but also with dissociative states such as following ketamine administrations and dreams, increased substantially in both regions (extended data Fig. 5 and 6). Compared to the naïve state, post-PCP GPe activity in post-mistake trials significantly increased, while the elevation of GPe activity in post-correct trials was found to be insignificant. These results correspond to the NHPs’ increased probability of initiating directed exploration and slightly lower probability of initiating random exploration compared to the naïve state (Fig. 2d,g). At the same time, the NHPs learned faster, making better predictions for finding the correct cues.

Finally, in the naïve state, the neural activity of both regions correlated with the learning slope, but the GPe also highly correlated with exploratory actions. Under PCP administration, DLPFC activity decoupled from the behavioral dynamics, and the NHPs’ ability to produce accurate predictions was reduced to chance levels. The GPe increased its activity, especially in post-correct trials, corresponding with the elevation in random exploration and contrary to the decrease in directed exploration. Post-PCP DLPFC activity correlated again with the learning slope, and the NHPs’ regained their ability to produce accurate predictions. The GPe increased its discharge rate further, especially during post-mistake trials corresponding with the increase in directed exploration and contrary to the decrease in random exploration. These findings (robustly found in separate NHP analysis, extended data Fig. 8), together with the increase in gamma activity in both regions post-PCP, led us to hypothesize that GPe activity is imperative for maintaining accurate exploratory strategies, perhaps by modulating attentional control over access to WM in the DLPFC through data dimensionality reduction^20^. We, therefore, propose that the increase in GPe activity under PCP administration was intensified by DLPFC dysfunction and impaired WM and predictive ability. The subsequent increase in GPe activity post-PCP, coupled with the restoration of predictive abilities, led to improved behavioral outcomes. Our results support the attention-gating hypothesis^19–21^, suggesting that the GPe plays a role beyond being a switch for exploration. The GPe may help to focus attention on relevant information and suppress irrelevant information, which is essential for accurate exploratory strategies.

### Adaptive reinforcement learning model

In search of a suitable test for our hypothesis, we developed an *adaptive Reinforcement learning (RL)* model that emulates the activity of the GPe by dynamically adjusting its learning gain parameter in response to changing learning demands (Fig. 3 and extended data Fig. 9). In our model, the simulated GPe (sGPe) activity (i.e., the learning gain parameter, α_t_) was adjusted based on environmental changes, proportional to the model’s knowledge of the task state and surprise level (Fig. 3a). The model’s predictive accuracy or greedy strategy was modified using a SoftMax probabilistic choice function, employing the Boltzmann distribution to generate predictions, wherein the ‘temperature’ parameter (T) controlled the action selection noise level (Fig. 3a).

**Figure 3:**
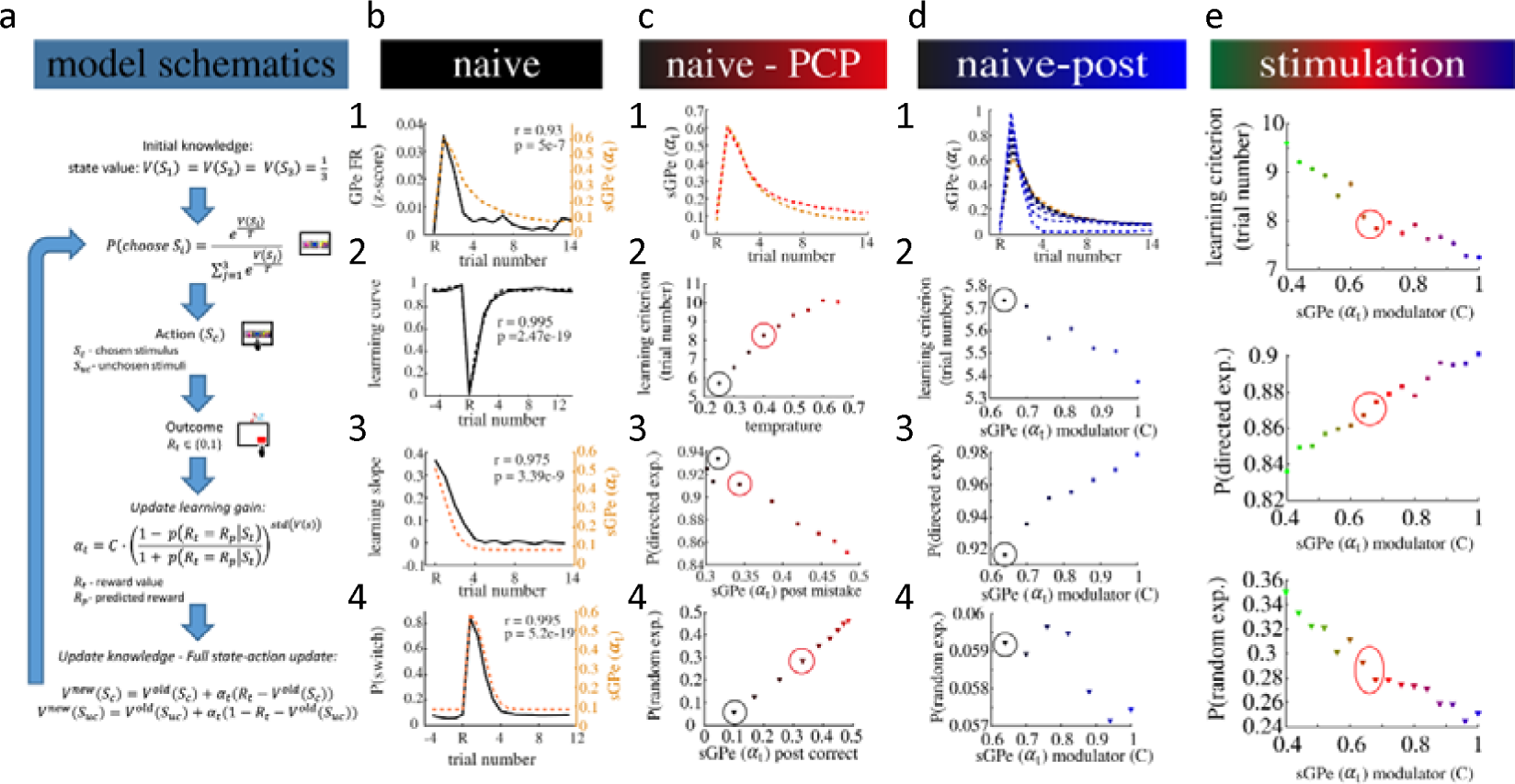
An adaptive reinforcement learning model replicates neuro-behavioral results and predicts potential benefits for GPe stimulation in NHPs modeled PCP-induced schizophrenia. **(a)** The model’s schematics, equations, and flow. **(b) Naïve –** simulating the model’s behavior and sGPe (a_t_) values best fitting the NHPs’ naïve results **(1)** Comparing the NHPs’ recorded GPe FR (solid black) with the model’s sGPe value (dashed yellow) calculated based on the NHPs’ choices (i.e., the model was not free to make its own choices. It was forced to replicate the NHPs’ choices and the sGPe value was calculated accordingly). **(2-4)** Comparing the model’s behavioral results and sGPe values to the NHPs’ data. **(2)** Comparing the simulation and the NHPs’ behavioral results. The NHPs’ (solid line) and the model’s (dashed line) learning curves. **(3-4)** Comparing the simulations learning dynamics with sGPe value. **(3)** The learning slope (solid black) and sGPe (dashed yellow). **(4)** switch probability (solid black) and sGPe value (dashed yellow). **(c) Naïve-PCP (1)** Comparing the model’s sGPe value calculated based on the NHPs’ naïve state (dashed black, as shown in naïve state figure 1) and PCP-state (dashed red) behavioral choices (i.e., calculating sGPe value according to the naïve and PCP state behavioral choices). **(2-4)** Simulating the PCP schizophrenia-modeled state by increasing the model’s temperature (i.e., reducing the model’s prediction accuracy). Simulated parameters color graded, changing the temperature from naïve values, T = 0.25 (black) to 0.65 (red). Black circles show the value under naïve parameters (i.e., values used for the model presented in the naïve state). The red circles indicate the ‘temperature’ value best fitting the NHP’s PCP behavioral results. **(2)** Learning criterion against temperature value. **(3-4)** Comparing the model’s change in exploratory behavior as temperature increases. **(3)** Directed exploration probability (i.e., the probability to switch choice following an unsuccessful trial) against sGPe value following a prediction-outcome mismatch as temperature increases from black to red. **(4)** Random exploration probability (i.e., the probability of switching following a successful trial) against sGPe value following a congruent prediction-outcome (i.e., following a successful trial) as temperature increases from black to red. **(d) Naïve-post: (1)** Comparing the model’s sGPe value calculated based on the NHPs’ naïve state (dashed yellow, as shown in naïve state figure 1) and post-PCP state (dashed black) behavioral choices (i.e., calculating sGPe value according to the naïve and post-PCP state behavioral choices). Then, simulating the increase in GPe’s activity observed post-PCP by increasing sGPe’s modulator parameter (C) from black to blue. **(2-4)** Simulating the increase in GPe’s activity observed post-PCP by increasing the modulator parameter (C) of sGPe and its effect on the model’s behavioral policy. Simulated parameters color graded, changing C_a_ parameter from baseline (naïve values, black, initial value indicated by a black circle) up (blue). **(2)** Learning criterion against sGPe modulator value. **(3-4)** Comparing the model’s change in exploratory behavior as sGPe modulator value increases. **(3)** Directed exploration probability (i.e., the probability of switching choice following an unsuccessful trial) against sGPe value following a prediction-outcome mismatch as sGPe modulator increases from black to blue. **(4)** Random exploration probability (i.e., the probability of switching following a successful trial) against sGPe value following a congruent prediction-outcome (i.e., following a successful trial) as sGPe modulator value increases from black to blue. **(e) Stimulation:** Simulating the effects of GPe stimulation under PCP conditions by increasing (graded blue) and decreasing (graded green) sGPe modulator parameter from values best fitting PCP results (red). **Top** – learning rate, **middle** – directed exploration probability and **bottom** – random exploration probability. Red circles indicate the values most fitting PCP results. **See also extended data figures 9.**

First, we tested whether the proposed sGPe acts similarly to the recorded GPe activity. We, therefore, ‘forced’ the model to replicate the NHPs’ behavioral choices (i.e., the model was ‘fed’ with the NHPs’ behavioral data) and calculated the resultant sGPe values. We found sGPe activity to be highly correlated with the NHPs’ recorded GPe discharge rate (Fig. 3b). The modeled sGPe is computed based on the previous reward outcome. This assumption is biologically viable and supported by the finding that GPe activity is susceptible to prediction-outcome mismatch (elevating its activity in response to a mismatch) and that this information is carried across trials (extended data Fig. 3c-d).

Next, we assessed whether the model could reproduce the observed neuro-behavioral outcomes when faced with the behavioral task (i.e., the model was now free to make its own choices to solve the task). Indeed, the model’s behavior showed a high degree of correlation with the NHPs’ behavioral data (Fig. 3b). Moreover, the model’s sGPe value was highly correlated with the learning slope and switch probability (Fig. 3b), as was the NHPs’ GPe activity (Fig. 1f). This suggests that the model’s sGPe value may play a similar role in learning and decision-making as the NHPs’ GPe activity.

We then wished to test our hypothesis that the increase in GPe activity under PCP administration was intensified by DLPFC dysfunction and impaired WM and predictive abilities. If our hypothesis is correct, and the increase in GPe activity is at least partially due to the increased uncertainty, then inputting the PCP-administered NHPs’ behavioral choices to the modeled sGPe should result in a similar change in its activity pattern. We, therefore, evaluated the simulated sGPe activity calculated according to the PCP-administered NHPs’ behavioral choices and compared it with the simulated naïve-state results calculated previously (Fig. 3c). Similar to PCP-induced GPe elevation post-correct (Fig. 2f, inset), the model’s exposure to PCP-state behavioral data raised later-block sGPe values (Fig. 3c).

To test the ability of our model to simulate the schizophrenia-modeled PCP results, we ‘freed’ the model to make its own choices while increasing its ‘temperature’ parameter (T) to create a noisier action-selection process as behaviorally observed. This change caused the model to learn slower (Fig. 3c), substantially increase random exploration, and slightly decrease directed exploration probabilities (Fig. 3c), which aligned with the behavioral changes observed in the schizophrenia-modeled NHPs. Furthermore, sGPe value modifications mirrored the recorded GPe activity changes, slightly elevating post-mistake (negatively correlated with directed exploration probability) and substantially elevating post-correct (positively correlated with random exploration probability) (Fig. 3c). These changes in sGPe values and exploratory policies in response to the elevated uncertainty replicated the observed alterations in the NHPs’ GPe discharge rates and exploratory behavior under PCP treatment (Fig. 2). The model also replicated the reduction in correlation between GPe activity and switch probability observed in the NHPs’ schizophrenia-modeled state (extended data Fig. 9a-b). These results reproduce the observed behavioral changes and replicate the concurrent alterations in GPe neural activity under PCP administration. Thus, a single parameter modification effectively produced the observed behavioral and neural alterations.

Subsequently, we investigated the model’s capacity to explain the neuro-behavioral changes observed following the cessation of PCP administration. Post-PCP administration, the DLPFC regained its correlation with the NHPs learning slope, and the GPe elevated its discharge rate further. Behaviorally, the NHPs’ predictive abilities were regained and even improved (Fig. 2), and their exploratory strategies sharpened. We first tested the ability of the improved behavioral strategy to impact sGPe values. Hence, we input the NHPs’ post-PCP choices into the naïve sGPe model to derive corresponding sGPe values. Comparison of sGPe values calculated from the NHPs’ post-PCP and naïve choices yielded similar outcomes (Fig. 3d). To simulate the recorded elevation in GPe discharge rate post-PCP, we increased the sGPe modulator value (’C’), thus increasing sGPe’s ‘baseline’ level, and recalculated sGPe values based on the same post-PCP behavioral choices (Fig. 3d). This resulted in an increase in sGPe activity, mimicking the observed change in GPe’s activity post-PCP (Fig. 3d).

Next, we tested whether this increase in baseline activity would also reproduce the observed behavioral results. We restored the temperature to its ‘naïve’ value (simulating the return of the NHPs’ predictive abilities) and systemically increased the sGPe modulator parameter (’C’) linearly (simulating the increase in GPe discharge rate) while ‘freeing’ the model to solve the task on its own. Consistent with the NHPs’ post-PCP results, increasing the sGPe modulator value enhanced the learning speed and directed exploration likelihood but reduced random exploration probability (Fig. 3d), mimicking the observed neuro-behavioral outcomes.

Finally, we explored whether varying the sGPe modulator parameter under heightened temperature (simulating PCP administration) could improve behavioral performance. Adjusting the temperature to mirror the NHPs’ PCP-induced results, we systematically varied the sGPe modulator parameter (Fig. 3e). We found that elevating the value of the sGPe modulator increased the model’s probability of engaging in directed exploration and decreased its likelihood for random exploration (Fig. 3e). This enhanced accuracy in exploratory strategies resulted in faster learning (Fig. 3e). Conversely, reducing the value of the sGPe modulator yielded the exact opposite effects (Fig. 3e). Thus, the adaptive RL model indicates that enhancing GPe activity may confer benefits for task performance in our NHPs with schizophrenia-like symptoms.

### GPe stimulation alters schizophrenia-modeled E-E balance

To test the prediction of the *adaptive RL* model and investigate the causal relationship between GPe activity and behavioral performance in the schizophrenia-modeled state, we conducted macro-electrode stimulation of the GPe during the NHPs’ behavioral task performance. We administered either high-frequency (130 Hz, activity dampener^33–35^) or low-frequency (13 Hz, activity enhancer^36–38^) stimulation to the GPe of the schizophrenia-modeled NHPs (Fig. 4a). Our results demonstrated that the frequency of stimulation had a significant impact on task performance, with 130 Hz stimulation leading to poorer performance and 13 Hz stimulation resulting in improvement (Fig. 4a,b).

**Figure 4:**
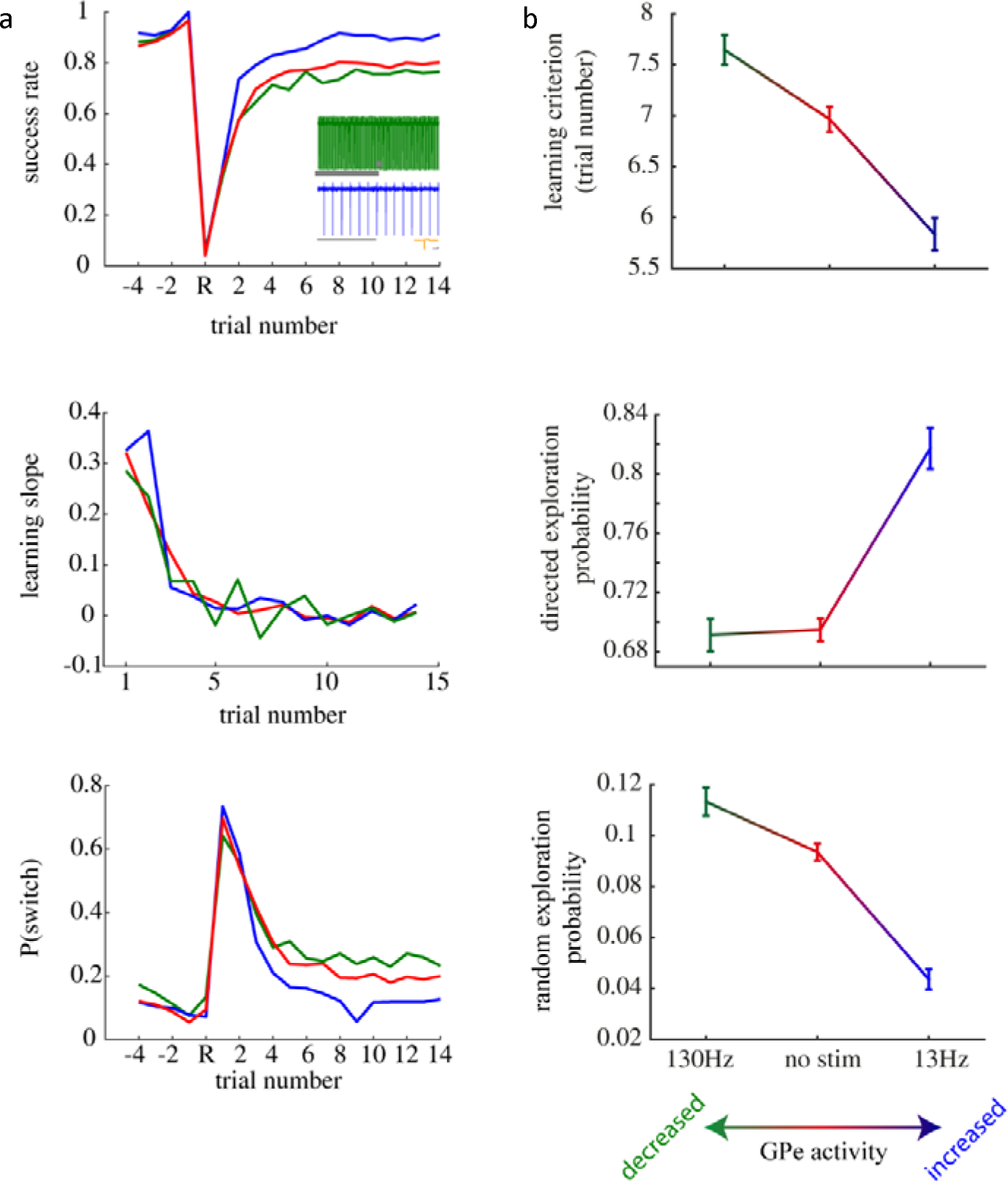
GPe 13 Hz macro stimulation improves task performance, whereas 130 Hz hampers it in the NHP PCP model of schizophrenia. (a) Behavioral performance analysis under PCP administration. Red - No stimulation, blue – 130 Hz continuous stimulation (activity dampener), and green – 13 Hz continuous stimulation (activity enhancer). **Top** – learning curve. Inset shows a one-second recording during 130 Hz stimulation (green) and during 13 Hz stimulation (blue), the grey bar shows 500 ms duration and a recording of one stimulation epoch (The horizontal grey bar shows 1 ms duration and the vertical line shows 1000 V). **Middle** – learning slope. **Bottom** – Switch probability. **(b) Top -** learning criterion (achieving three consecutive correct choices) during 130 Hz stimulation (left, blue), no stimulation (middle, red) and 13 Hz stimulation (right, green). **Middle -** The NHPs probability of making directed exploration (i.e., to switch their choice after a prediction-outcome mismatch). **Bottom** - The NHPs’ probability of making random exploration (i.e., to switch their choice after congruent prediction-outcome). **See also extended data figures 10.**

Decreasing GPe activity reduced the probability of directed exploration while increasing its activity enhanced it (Fig. 4b). Furthermore, the NHPs’ learning dynamics improved when GPe activity was enhanced, leading to faster learning and improved learning slope (Fig. 4a). These changes in the directed exploration phase of the task translated into slower learning when GPe activity was decreased and faster learning when it was increased (Fig. 4b). Furthermore, decreasing GPe activity increased random exploration, whereas increasing its activity decreased it (Fig. 4b). These robust results were consistent across both NHPs (extended data Fig. 10).

The stimulation behavioral effects highlight the modulatory role of GPe activity in directed and random exploration abilities within the schizophrenia-modeled NHPs. As GPe activity intensified, exploration patterns became more accurate, leading to an increased probability of initiating directed exploration and a decreased likelihood of performing random exploration. Conversely, reducing GPe activity resulted in less accurate exploration patterns. In general, GPe activity’s opposite effect on directed and random exploration probabilities suggests that GPe serves a function in controlling E-E balance beyond being an exploration generator^27^.

## Discussion

Maintaining the E-E balance is imperative for learning; hence its neural underpinnings have long been debated. The DLPFC, the center of executive functions, and the GPe, through GPe-subthalamic and other BG main-axis loops, have both been suggested as the primary hubs controlling this balance^14–18,39^. Alternatively, GPe’s primary role is to dynamically regulate WM, stored in cortical areas, according to the changing environment and the agent’s needs^19–21^. This gating is probably maintained by reducing the dimensionality of the information sent from the whole cortex to the BG and back again to the frontal cortex^15,20^.

Schizophrenia patients have been observed to have dysfunctional cortical and GPe activity^23–26^. They also manifest behavioral E-E imbalance, often exhibiting excessive random exploration strategies, prioritizing them over directed exploration^1,2^. Therefore, the E-E balance has recently been suggested to be a holistic and ecologically valid framework to resolve some of the apparent paradoxes that have emerged within schizophrenia research^2^. Here we show that increased GPe activity improves exploratory strategies (both directed and random) and propose an *adaptive RL* model replicating the observed results. Our experimental and model results suggest that the GPe acts as a dimensionality reduction mechanism to increase ‘attention’ to prominent features and controls E-E balance indirectly. This hypothesis is supported by the BG anatomical funneling structure (the cortico-basal ganglia-thalamo cortical network converges to the globus pallidus, and neuronal numbers decline through striatum, GPe, GPi) and recent model and human findings^15^.

Attention is proposed to act as a ‘gatekeeper’ for WM^19,40^, and both processes are associated with gamma oscillatory activity^41^. In schizophrenia, gamma activity alterations have been suggested as a biomarker for glutamatergic dysfunction^32^. Consistent with this finding, NMDA-R antagonists have been found to increase gamma activity and synchrony in the GPe^42^, further reducing the dimensionality of the information funneled through the basal ganglia and back to the cortex. The increase in synchrony may either be a compensatory mechanism or a primary defect impairing cognitive and WM abilities. Our results extend the understanding of the role of GPe gamma activity, showing that its increased activity correlates with increased GPe activity and improved exploration patterns and WM performance in the post-PCP period. In the future, it will be interesting to explore gamma activity-specific parameters as more accurate schizophrenia biomarkers similar to those suggested in adaptive DBS for Parkinson’s disease^43^ and obsessive-compulsive disorder^44^ (e.g., synchrony and coherence with other activity bands, most notably theta).

In the current task framework, the relationship between WM and the prediction of correct choice is evident. Task design demands using WM as predictions (prior) in the subsequent trial. It is, therefore, enticing to discuss our results through the predictive coding prism. Predictive coding theory hypothesizes that the cause of psychosis is reduced precision of prior beliefs and/or increased accuracy of sensory data (likelihood) and that NMDA-R dysfunction may impair their balance^45^. Here we show, using the NMDA-R antagonist model of schizophrenia, that increasing GPe activity using low-frequency macro-stimulation results in better, more accurate predictions, minimizing the gap between prior beliefs and likelihood. We, therefore, hypothesize that GPe low-frequency DBS may be advantageous for schizophrenia management.

Schizophrenia is a significant health burden estimated to affect 21 million people globally (0.3-0.5% of the adult population)^46^. Unfortunately, up to 34% of schizophrenia patients are treatment-resistant^47^ and are offered limited treatment options after first-line therapies have failed, with many patients continuing to have symptoms despite maximal therapy. For these reasons, recent research has focused on finding an appropriate site for DBS in schizophrenia patients. The suggested areas for stimulation are the nucleus accumbens, hippocampus, globus pallidum – internal segment (GPi), dorsomedial thalamus, and medial septal nucleus^48^. Recent studies showed that the lamina between GPe and GPi might be the optimal target for DBS treatment of Parkinson’s disease^49^. Similarly, our results support targeting the GPe for DBS treatment in schizophrenia.

## Data availability

The dataset supporting the current work is available from the corresponding author upon request.

## Code availability

The analysis code and supporting the current work will be made publicly available on GitHub.

## Contributions

N.A conceived the research, designed the experiments, performed the in vivo experiments (electrophysiological and behavioral recordings) and the surgical procedures, performed data analysis, statistics, and mathematical modeling. N.A, N.R and U.W-R performed the in vivo experiments. NA and Z.I performed the surgical procedure. H.B conceived the research, designed the experiments, and supervised the work. N.A wrote the manuscript with input from H.B. All authors critically reviewed the manuscript.

Correspondence should be sent to Nir Asch.

## Materials and Methods

### Animal Training and Behavioral Tasks

Data were obtained from two female vervet monkeys (Cercopithecus aethiops, monkeys K and R) weighing 3.5–4 kg. All data were pooled for both monkeys for all analyses. Care and surgical procedures were in accordance with the National Research Council Guide for the Care and Use of Laboratory Animals^1^ and the Hebrew University guidelines for the use and care of laboratory animals in research, supervised by the institutional animal care and use committee of the Hebrew University and Hadassah Medical Center. The Hebrew University is an Association for Assessment and Accreditation of Laboratory Animal Care internationally accredited institute. The behavioral paradigm used was a multiblock three-choice reversal learning task (Fig. 1 b). The NHPs used their right (contralateral to the recording side, discussed below) hands to touch stimuli presented on a screen that was located ∼16 cm from their heads (Elo 1939L 19-inch open-frame touch-monitor; Elo Touch Solutions Limited).

In each daily session, three square fractal images were randomly selected as stimuli out of a set of 10 possible stimuli. The stimuli were presented in fixed positions throughout the session, on the left, right, and center of the white screen. Each trial began with a presentation of a black horizontal box on the lower right corner of the screen (Fig. 1, b). To initiate a trial, the NHPs touched the black rectangle, which then disappeared. Two seconds later, the three fractal stimuli appeared. After touching (choosing) one of the three, all three stimuli disappeared. Two seconds later, a red horizontal box appeared on the bottom of the screen. Touching this box was followed by three simultaneous results: The box would disappear, a banana-flavored liquid reward was either delivered or not (according to the monkey’s choice, discussed in the next paragraph), and an ∼80ms auditory stimulus was played, obscuring any acoustic artifacts of the food pump. The sound played independently of the trial’s outcome. Attempts were aborted if no choice was made within 30 seconds or the red box was not touched within 30 seconds. All trials (correct, incorrect, and aborted) were followed by a variable intertrial interval (ITI) lasting 5–8s. For each block, only one of the three stimuli was deterministically rewarded. No reward was delivered for the choice of other stimuli. The monkeys had no guiding information pointing them toward the correct stimulus, but eventually, learning was established, and the probability of choosing the correct stimulus reached a plateau (Fig. 1c). The criterion for learning was reached once the monkey chose the rewarded stimulus for 12–15 trials out of the last 25 (the criterion was randomly selected per block). Once this happened, an un-cued switch in the reward stimulus’s identity occurred (i.e., reversal), and a new block started. Each daily session ended once the monkeys no longer initiated trials. After long periods in which the monkeys did not work, the experimenter occasionally delivered a free reward to remotivate them. The behavioral paradigm was designed and run using the Psychophysics toolbox (Brainard et al., 1997) for MATLAB (The MathWorks, Inc.). Monkeys were trained for 5–6 days per week and were allowed free access to water in their home cages. Supplementary food was delivered when the monkeys did not reach the predefined daily calorie minimum. Monkeys were given free access to food on the weekends.

### Surgery and MRI

The monkeys were fully trained on the task (4–5 months) before the recording chamber was implanted (Fig. 1a). After the training period, they were operated on under full anesthesia and sterile conditions. In the surgery, an MRI-compatible Cilux head holder (Crist Instrument) and a square Cilux recording chamber (AlphaOmega) with a 27mm (inner) side, located above a burr hole in the skull, were attached to the heads of the monkeys. The recording chamber was attached to the skull, tilted ∼45° laterally in the coronal plane, with its center targeted at the stereotaxic coordinates of the left GPe. All surgical procedures were performed under aseptic conditions and general isoflurane and N2O deep anesthesia. Analgesia and antibiotics were administered during surgery and continued postoperatively. Recording began after a postoperative recovery period of several days. We estimated the stereotaxic coordinates of the recording target using MRI scans. The MRI scan (General Electric 3 tesla system, T2 sequence) was performed under i.m. Domitor and ketamine sedation. Upon experiment completion, all surgical attachments were removed from NHPs. The NHPs were then rehabilitated and placed at the Israeli Primate Sanctuary.

### Recording and Data Acquisition

During recording sessions, the heads of the monkeys were immobilized, and up to eight glass-coated tungsten micro-electrodes (impedance 0.15–1MΩ at 1,000 Hz), confined within a cylindrical guide (1.65-mm inner diameter) were advanced separately (EPS; Alpha-Omega Engineering) into the GPe and the DLPFC. The electrical activity was amplified with a gain of 20, then filtered using hardware Butterworth filters (high-passed at 0.075 Hz, two poles; low-passed at 10,000 Hz, three poles) and finally sampled at 44.6 kHz (SnR; Alpha-Omega Engineering). Cortical and GPe units were identified by the stereotaxic and MRI coordinates, by the electrophysiological hallmarks of the encountered structures along the penetration, and by their own unique characteristics, such as GPe neurons’ high FRs and pausing behavior^2^. Neuronal activity was sorted and classified online using a template-matching algorithm (SnR; Alpha-Omega Engineering). Each position of entry was registered according to the chamber’s X-Y coordinates, and brain structures were identified electro-physiologically. Cells were selected for recording as a function of their isolation quality and optimal signal-to-noise ratio^3^. Thus, creating a 3D map of the desired brain column beneath the chamber.

### Analysis of Behavior

To evaluate the NHPs’ exploratory behavior, we categorically divided exploration patterns into exploration-favorable (following a prediction-outcome mismatch) and exploitation-favorable (without such mismatch) trials. The former are trials that followed an unsuccessful trial (i.e., post-mistake) in which exploration (i.e., switch of choice) will lead to reward, and the latter are trials that followed a successful trial (i.e., post-correct) in which exploitation (i.e., choosing the same stimulus as before) will lead to reward. Each category was further subdivided into two groups, resulting in four options. (1) Directed exploration – trials following an unsuccessful trial in which the NHPs’ switched their choice (i.e., explored). (2) Perseveration – trials following an unsuccessful trial in which the NHPs’ did not switch their choice. (3) Random exploration – trials following a successful trial in which the NHPs’ switched their choice. (4) exploitation –trials following a successful trial in which the NHPs’ did not switch their choice.

We also evaluated the learning dynamics throughout the task, looking at five main parameters. The learning curve, defined as the NHPs’ success rate (i.e., the probability of choosing successfully), the learning slope (the derivative to the learning curve), switch probability (i.e., the probability of choosing a different cue than the choice of the previous trial), which was further divided to successful switch probability (i.e., the probability to switch successfully – finding the correct cue) and unsuccessful switch probability (i.e., the probability of switching unsuccessfully). We further analyzed the NHPs’ probability of success if they switched their choice (i.e., the probability of success given that a switch was made). This analysis provides more precise information about the NHPs’ memory of the previous states (50% chance is a random choice).

Lastly, we evaluated the NHPs’ response times. Response times were defined as the length of time it took the NHPs to 1) choose the presentation of the three stimuli and 2) claim the reward outcome by pressing the red rectangle (i.e., reaction plus movement time). All data analysis was performed using MATLAB (The MathWorks, Inc.).

### Analysis of Single Unit Firing Activity

Data analysis was conducted only for units that were stably held, well-isolated [isolation score (24) ≥0.8], and unquestionably identified as either cortical or GPe units. Only GPe units that fired ≥20 spikes per second on average were included (thus focusing on the prototypical population). Continuous traces showing GPe spiking activity (Fig. 1a) were obtained using a two-pole Butterworth filter (1,000–3,000 Hz). Background neural activity was recorded during the last three seconds of the inter-trial interval. We defined a baseline period of two seconds for calculating the change from resting activity for the plots showing the z-scored dynamic temporal patterns of GPe and DLPFC activities (i.e., post-stimulus time histograms, PSTH). We divided each second into ten nonoverlapping bins and calculated the spike count per bin. This data, per trial, was used as a baseline for the PSTHs. For each time point of the PSTH, the baseline discharge rate for that trial was subtracted from the count per bin, and the difference was divided by the baseline SD. We used the last two seconds of the ITI as the baseline for all epochs. The obtained values were filtered in all cases using a 50ms-wide Gaussian Kernel (MATLAB’s filtfilt function).

### Chronic, low-dose PCP administration

A mini-osmotic pump (Alzet osmotic pump model 2ML4) was implanted subcutaneously between the scapulae of both NHPs, providing a continuous daily dose of 1.68 mg/kg for 28 days. Both NHPs were implanted twice with at least one month washout period in between. Post-PCP recordings started at least two weeks after pump removals.

### Macro stimulation

Using the 3D map constructed during our recording sessions, we chose a desired stimulation position, maximizing the GPe location. We then placed a microgrid attached to the chamber’s inner wall. Using a specially designed guiding tower (AlphaOmega) we inserted a (recording) micro-electrode to the desired location, marked the GPe borders, and identified its center. Placing the concentric macro-electrode (microProbes CEAX 200) into the head mount allowed us to insert it into the desired location and stabilize it to the grid using dental cement. The macro-electrode remained connected and immobilized to the chamber. No sooner than three days after implantation, experimentation resumed. The NHPs were given either 13Hz or 130Hz macro-stimulation, on alternating days, during the behavioral task while we recorded their performance.

### Data analysis

Statistical analyses were performed using MATLAB version R2022b and are described in the figure legends. The correlation coefficient is represented by the letter ‘r’ for correlation analyses. The corresponding probability value, P, is the probability of getting ‘r’ as large as the observed value by random chance when the true correlation is zero and was computed by transforming the correlation to create a t-statistic. When there were multiple comparisons, the Bonferroni correction was used (i.e., the obtained P-value was multiplied by the number of comparisons) before significance testing (P<0.05) and is presented as such. All data presented met the assumptions of the statistical test used.

### RL model

Based on our neural and behavioral results, we hypothesized that the GPe acts as a gateway between automaticity and intentional behavior, exerting attentional control over access to working memory, necessary for effective exploration, to modulate the new state values (or predictions) in the DLPFC. Therefore, when the knowledge of the current state is known, the DLPFC can “rest” or be preoccupied with other tasks, but when the current state is unknown, the DLPFC is recruited by the GPe to modulate its knowledge and create new predictions (see the model’s schematics in Fig. 3).

To test our hypothesis, we created an *adaptive RL* model wherein the step-size parameter α changes its value according to the changing need for learning, adjusting its value according to the changing environment proportional to the model’s “surprise” and knowledge of the current task state. This model learns the value of choosing each stimulus available in the experiment using standard temporal difference learning. However, since the NHPs were highly proficient at performing the task and switched their choices rapidly after mistakes, we used a full state-action update role such that after choosing a stimulus *S*, and observing the reward for that choice *R_t_*, the value of all stimuli *V(S)* is updated according to the following:

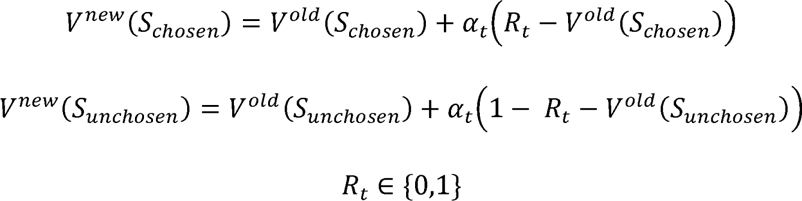

The learning rate parameter α controls the modulation of the stimulus values and hence the learning pace. Therefore, higher α values would correspond with faster, more vigorous learning (faster changing of the memories/predictions), and lower values would correspond with slower, potentially no learning.

To select one of the three stimuli available in each trial (*S*_1_, *S*_2_, *S*_3_), their current values are entered into a softmax probabilistic choice function, using the Boltzmann distribution to create predictions:

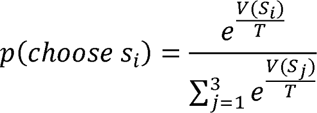

Such that the ‘temperature’ T sets the level of noise in the decision process, with large T corresponding to high decision noise with nearly random decision and small T corresponding to low decision noise and near-deterministic (greedy) choice of the highest-value option.

We hypothesized that the GPe (accepting the prediction error values [PE] encoded by dopamine projected to the striatum) modulates the learning rate in the DLPFC. Therefore, when PE value is high (i.e., the surprise of the current perceived reward is high) α should increase and allow fast learning. On the other hand, when PE is low (i.e., the surprise is low, and prediction of the current choice reward is accurate) α should decrease and no learning is needed.

We, therefore, defined 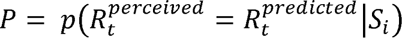 as the model assigned probability of receiving the perceived reward value 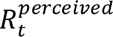 (we assumed negligible noise in the process of perceiving reward). For example, assume that *p*(*choose s*_1_) = 0.98 and *S*_1_was chosen. If 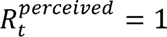, meaning reward was given then *P* = *p*(*choose s*_1_) = 0.98. But if reward was not given, then *P*=1 – *p*(*choose s*_1_) = *p*(*choose s*_2_) + *p*(*choose s*_3_) = 0.02.

We then used Shannon entropy to evaluate surprise, given a choice, and the perceived reward, -log (*P*). To normalize the surprise value to the range [0,1] and use it to modulate the value of the step-size parameter α, we will use the following function *f_surprise_* (*x*):

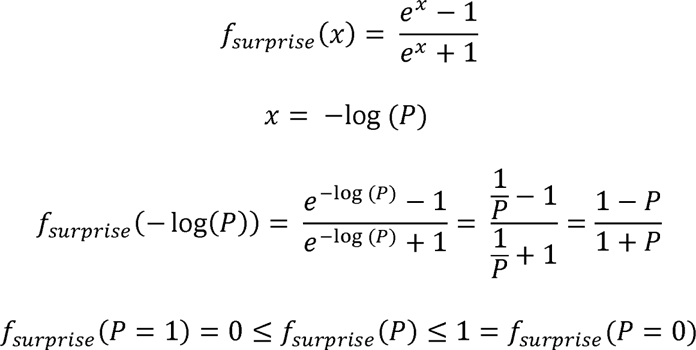

Therefore, as the mismatch between the prediction and the outcome is small (*P* → 1) surprise is low, and the learning rate decreases. But when the mismatch between the prediction and the outcome is large (*P* → 0) the surprise is high, and the learning rate increases.

To increase learning in situations of least knowledge (i.e., *V*(*S*_1_)= *V*(*S*_2_)= *V*(*S*_3_)) we raised *f_surprise_* by the power of the standard deviation (SD) of the value of their corresponding probabilities, *std*(*p*(*choose s*_1_), *p*(*choose s*_2_), *p*(*choose s*_3_)). Such that when the SD is low a will increase and when it is high it will not. Finally receiving the following update rule for a:

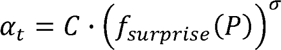

Where *C* is a constant, *C* ∊ (0,1) and σ is the SD of choice probabilities.

