## Supplementary figures for "Basal Ganglia Stimulation Ameliorates Schizophrenia Exploration Anomalies"

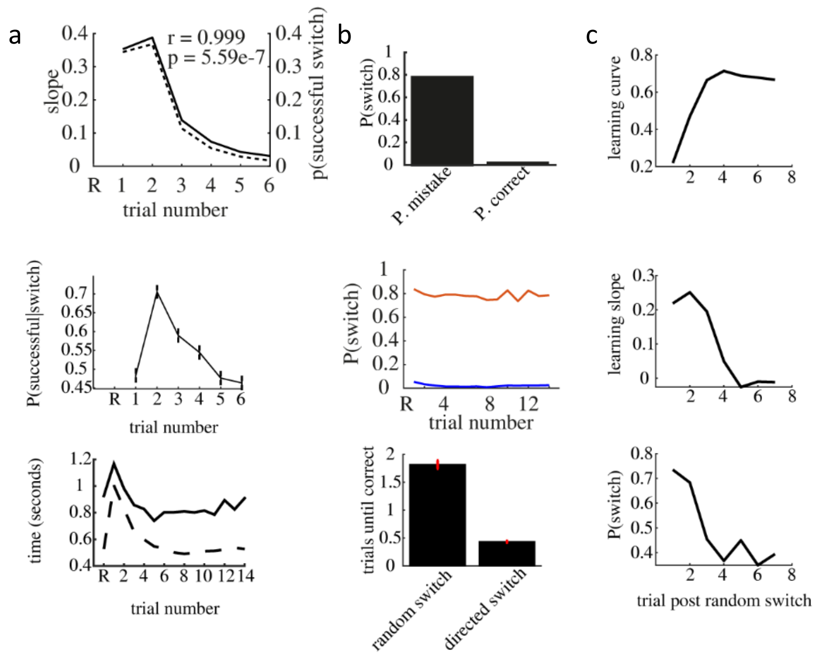


**Figure 1: Naïve state behavioral analysis. (a) The NHPs' learning dynamics. Top –** successful switch probability (i.e., the NHPs' probability of switching to the correct stimulus, solid line) and learning slope (dashed line), r and p represent the correlation between the two parameters and its corresponding P-value. **Middle** – the NHPs' probability of choosing successfully given that a switch was made (chance level is 0.5). **Bottom -** response time. Solid line - stimulus selection time, which is the duration between the appearance of stimulus choices (i.e., three fractal shapes) on the screen and the moment of response (by touching one of the fractal shapes). Dashed line – time to claim reward, from stimulus outcome appearance (red rectangle) until pressed to claim reward. **(b) Top -** Probability of response switching (i.e., choosing a different stimulus than the previous trial) after an unsuccessful trial (post-mistake) and after a successful trial (post-correct). **Middle -** the same as figure **d** according to trial number. Post mistake (red line) and post-correct (blue line) **Bottom -** the average number of trials needed until finding the correct response after random exploration (i.e., a switch following a successful trial) and after directed exploration (i.e., a switch following an unsuccessful trial). **(c)** Learning dynamics after random exploration.

**Related to main text figure 1.**

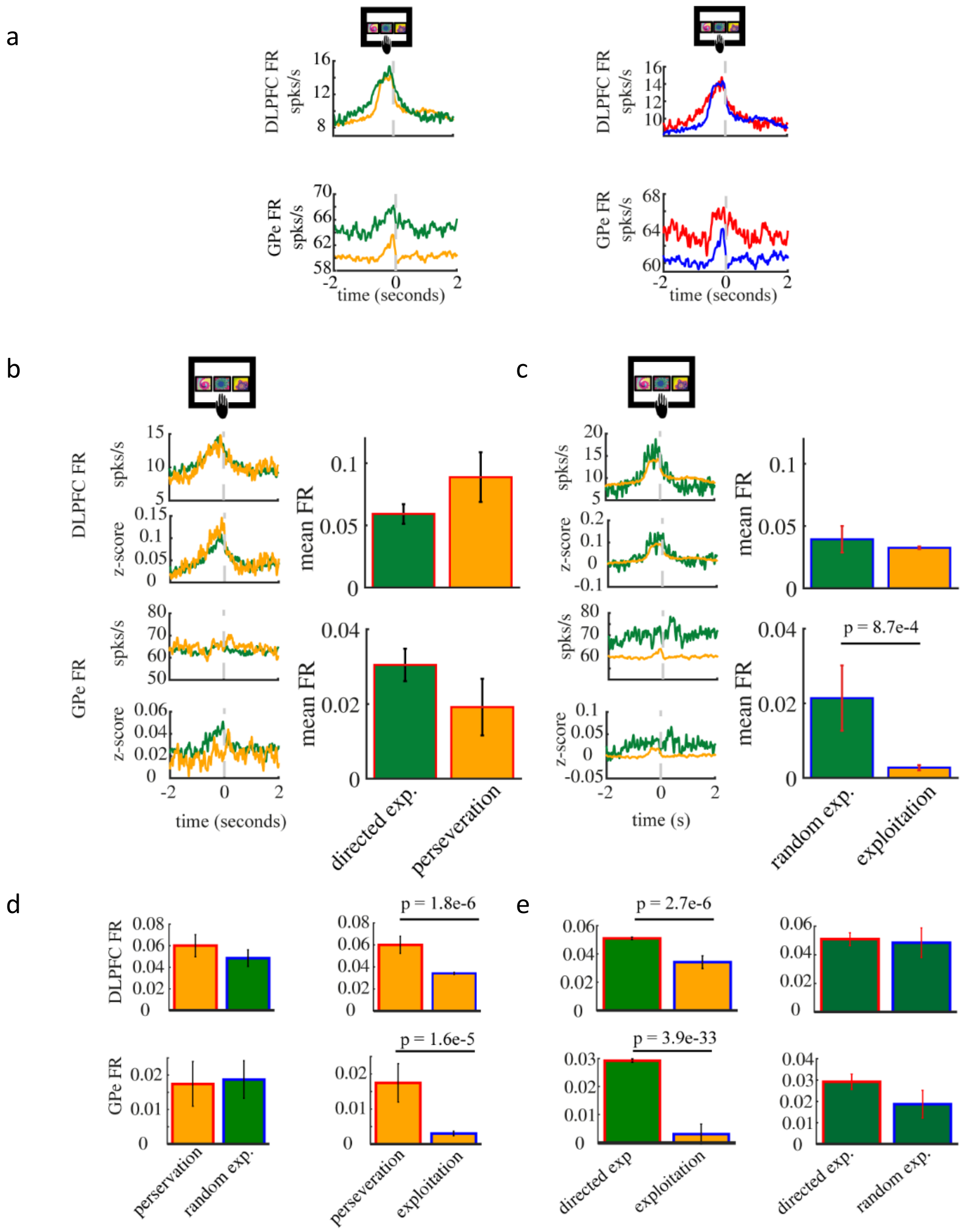


**Figure 2: Comparison of DLPFC and GPe firing patterens around choice selection. (a) Top left -** DLPFC firing rate around choice selection comparing switched and same choice trials. **Bottom left** – the same for the GPe. **Top right -** DLPFC firing rate around choice selection, comparing post-mistake, red and post-correct, blue. **Bottom right** – the same for the GPe. **(b)** Comparing DLPFC and GPe FR around choice selection in directed exploration (switched post mistake) and perseveration (same choice post mistake) trials. Bar graphs represent the mean and SEM FR during the two seconds preceding stimulus choice selection. Ps represent the p-value of the two-sampled t-test comparing the mean FR values. P-values are Bonferroni corrected (i.e., p-values are multiplied by the number of multiple comparisons). This annotation follows in all subsequent plot. **(c)** Comparing DLPFC and GPe FR around choice selection in random exploration (switched post correct) and exploitation (same choice post correct) trials. Bar graphs represent the mean and SEM FR during the two seconds preceding stimulus choice selection. **(d)** Comparing DLPFC and GPe FR around choice selection between perseveration trials (same choice post mistake) and random exploration (left) and exploitation (right) trials. **(e)** Comparing DLPFC and GPe FR around

choice selection between directed exploration trials (switched post mistake) and exploitation (left) and random exploration (right) trials.

**Related to main text figure 1.**

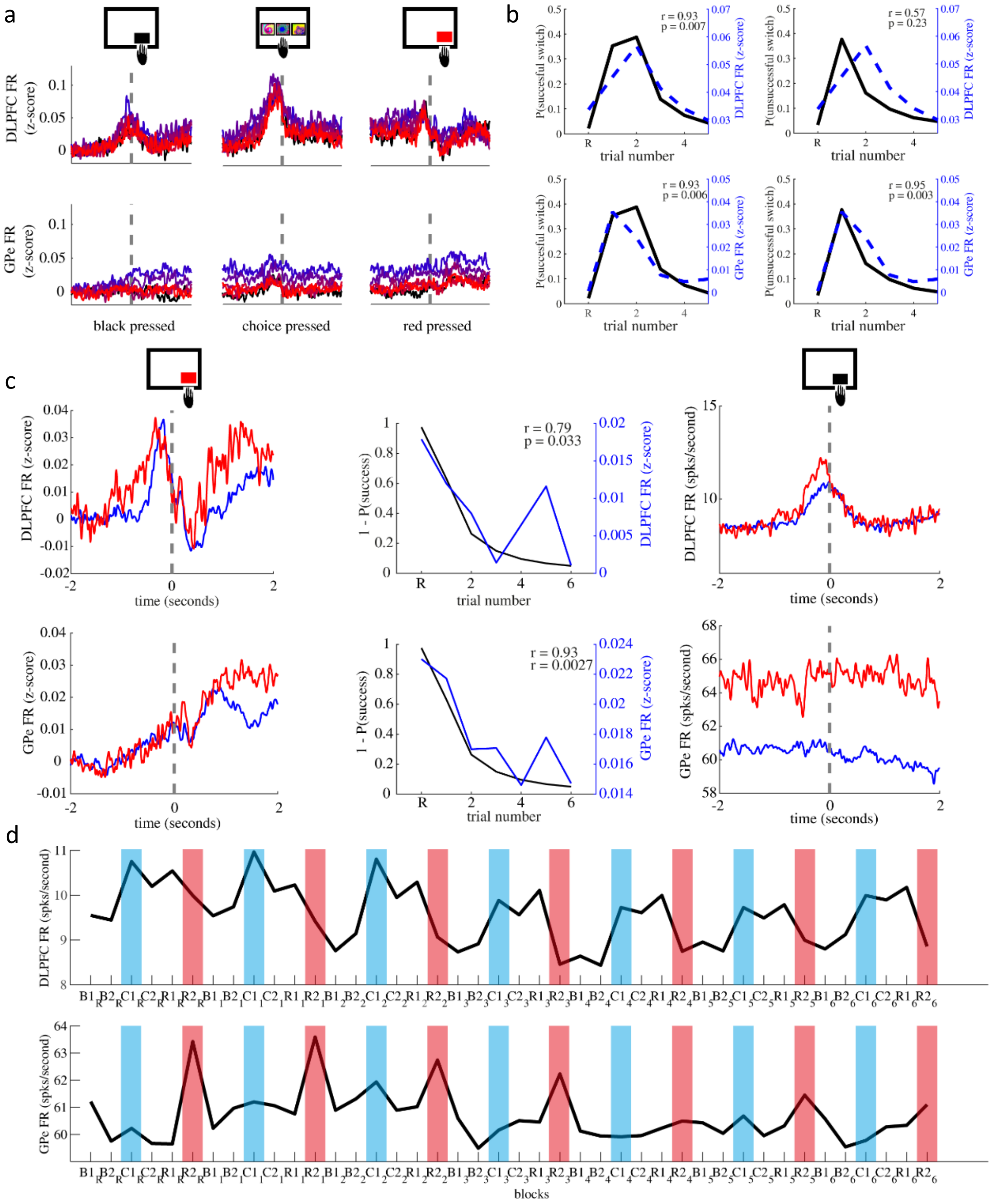


**Figure 3: DLPFC activity is highest around choice selection and correlates with the probability of making a successful switch whereas GPe activity is highest around reward outcome, carrying the information to the subsequent trial. (a)** DLPFC (top) and GPe (bottom) neuronal discharge rates during task performance color coded from blue (early trials) to red (later trials). Reversal trial is colored in black. **(b) Left -** Mean, DLPFC (top) and GPe (bottom), firing rates of the two seconds preceding choice selection (dashed blue) with the NHPs' probability of making a successful switch (solid black). **Right -** Mean, DLPFC (top) and GPe (bottom), firing rates of the two seconds preceding choice selection with the

NHPs' probability of making an unsuccessful switch (solid black). **(c) Left –** DLPFC (top) and GPe (bottom) mean FR around stimulus reward outcome press (at time zero) z-scored by the FR's mean and standard deviation during the two seconds prior to stimulus choice selection to isolate the effect of reward on the regions' activity. Red represents unsuccessful trials' average and blue successful trials' mean. **Middle –** Comparing the mean FR during the two seconds after stimulus reward press with the probability of being unsuccessful. **Right –** DLPFC (top) and GPe (bottom) mean FR around trial initiation cue apperance (at time zero). Red represents trials following an unsuccessful trial and blue represents trials following a successful trial. **(d)** DLPFC (top) and GPe (bottom) mean FR with task progression from reversal trial (R) to the sixth trial. B1 – mean FR during the two seconds leading to trial initiation cue press (pressing the black box). B2 – mean FR during the two seconds that follow black box press. C1 – mean FR during the two seconds leading stimulus choice press. C2 – mean FR during the two seconds folloing stimulus choice press. R1 – mean FR during the two seconds leading to reward outcome press (pressing the red box). R2 – mean FR following reward outcome press.

**Related to main text figure 1.**

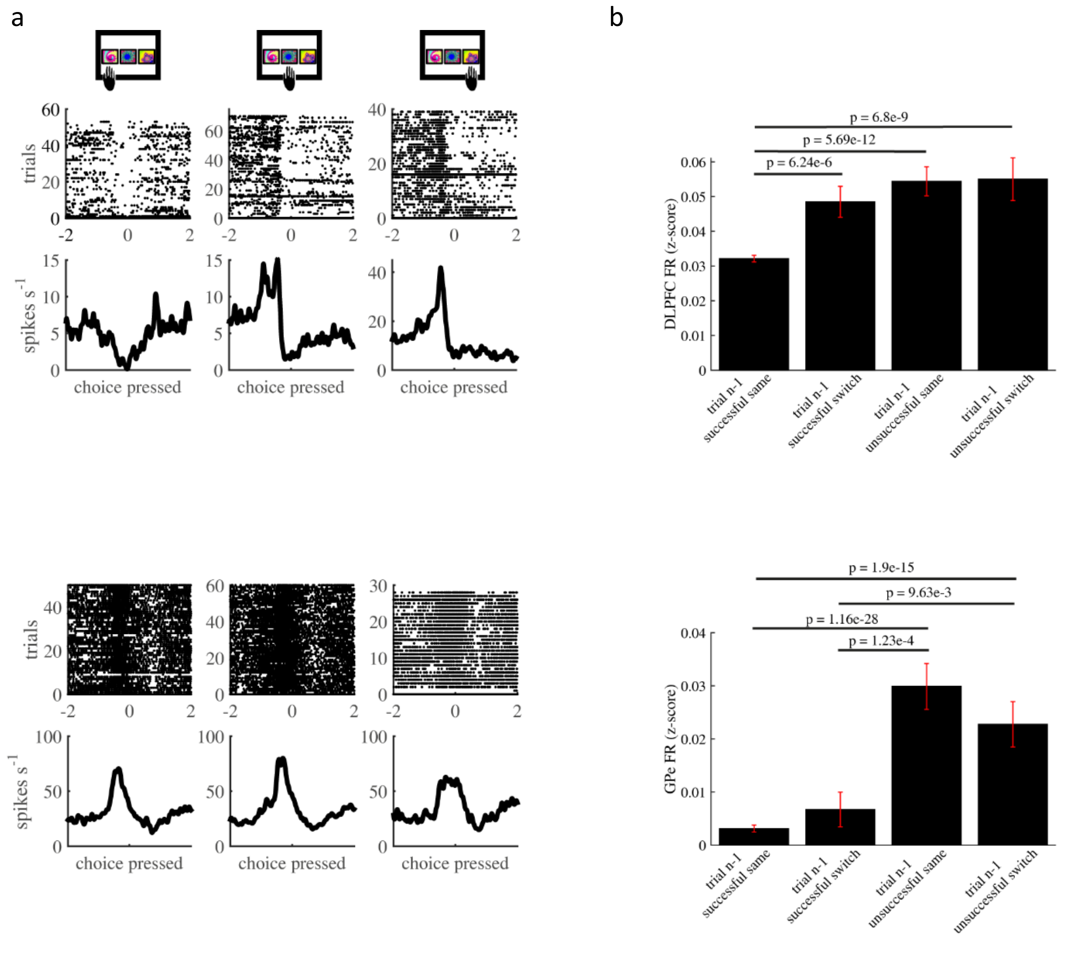


**Figure 4: DLPFC activity is more "sophisticated" and persists post-identification of a successful cue. (a) Top -** A PSTH of a single examplary DLPFC cell around stimulus choice selection of all three options. **Bottom** – A PSTH of a single examplary GPe cell around stimulus choice selection of all three options. **(b) Top –** mean DLPFC firing rate around choice selection following successful trials (switched and not switched) and following unsuccessful trials (switched and not switched). **Bottom -** mean GPe firing rate around choice selection following successful trials (switched and not switched) and following unsuccessful trials (switched and not switched).

**Related to main text figure 1.**

#### Extended data figures

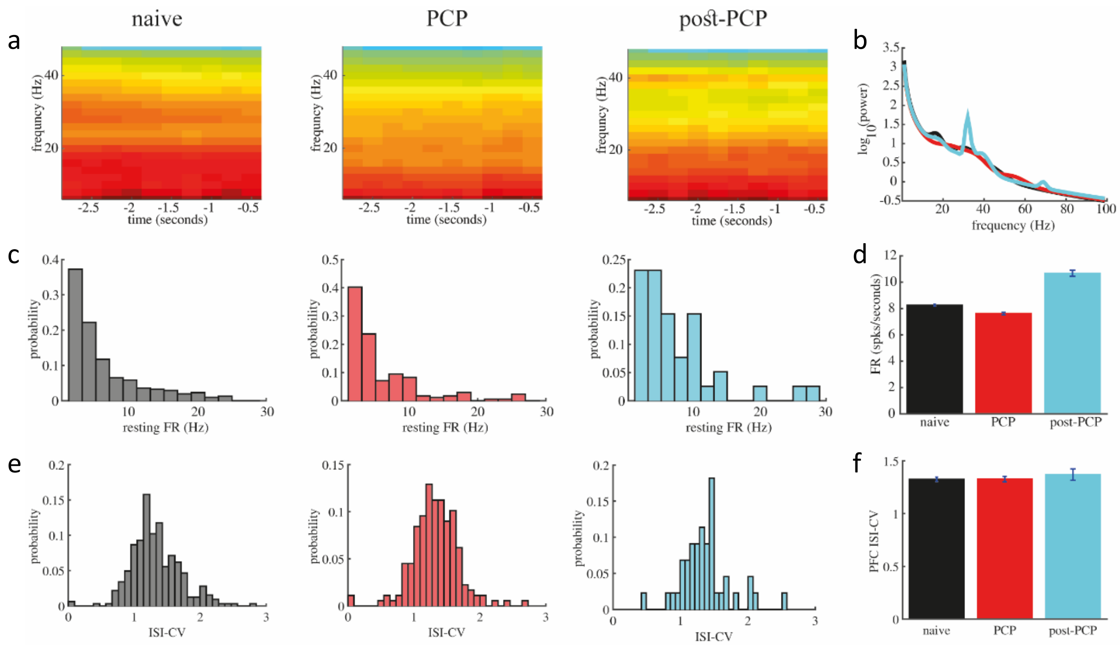


**Extended data figure 5: Background DLPFC neuronal changes under PCP adminstration and its long term effects. (a)** Spectrogram of DLPFC activity recorded during the three seconds preceding trial start. **(b)** Power spectrum of DLPFC neural activity recorded over the three seconds preceding trial start of the naïve state (black), under PCP administration (red), and during the post-PCP period (magenta). **(c)** Histogram of the mean resting state firing rate of the DLPFC neuronal cell population calculated during the three seconds preceding trial start. **(d)** Average firing rate and SEM of DLPFC activity during the three seconds preceding trial start. **(e)** Histogram of the inter-spike interval coefficient of variation of the DLPFC neuronal cell population calculated during the three seconds preceding trial start. **(f)** Average inter-spike interval coefficient of variation and SEM of the DLPFC activity recorded during the three seconds preceding trial start.

**Related to main text figure 2.**

#### Extended data figures

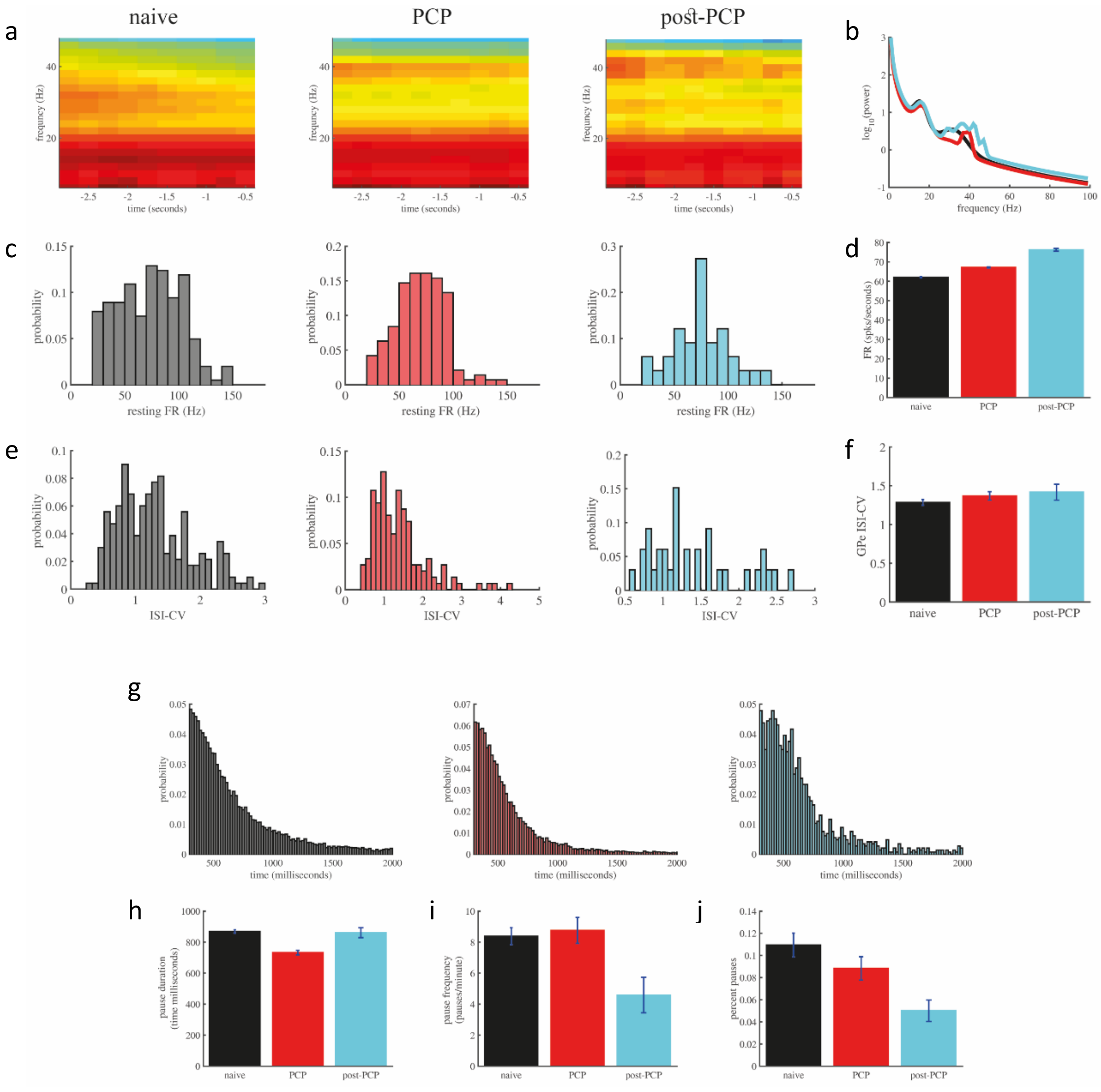


**Extended data figure 6: Background GPe neuronal changes under PCP adminstration and its long term effects. (a)** Spectrogram of GPe activity recorded during the three seconds preceding trial start. **(b)** Power spectrum of GPe neural activity recorded over the three seconds preceding trial start of the naïve state (black), under PCP administration (red), and during the post-PCP period (magenta). **(c)** Histogram of the mean resting state firing rate of GPe neuronal cell population calculated during the three seconds preceding trial start. **(d)** Average firing rate and SEM of GPe activity during the three seconds preceding trial start. **(e)** Histogram of the inter-spike interval coefficient of variation of GPe neuronal cell population calculated during the three seconds preceding trial start. **(f)** Average inter-spike interval coefficient of variation and SEM of the GPe activity recorded during the three seconds preceding trial start. **(g)** Histogram of the mean pause duration of GPe cells calculated over the entire stable recording time. **(h)** Average pause duration in all three conditions. **(i)** Average (and SEM) pause frequency. **(j)** Average (and SEM) percent of time in which GPe neurons pause.

**Related to main text figure 2.**

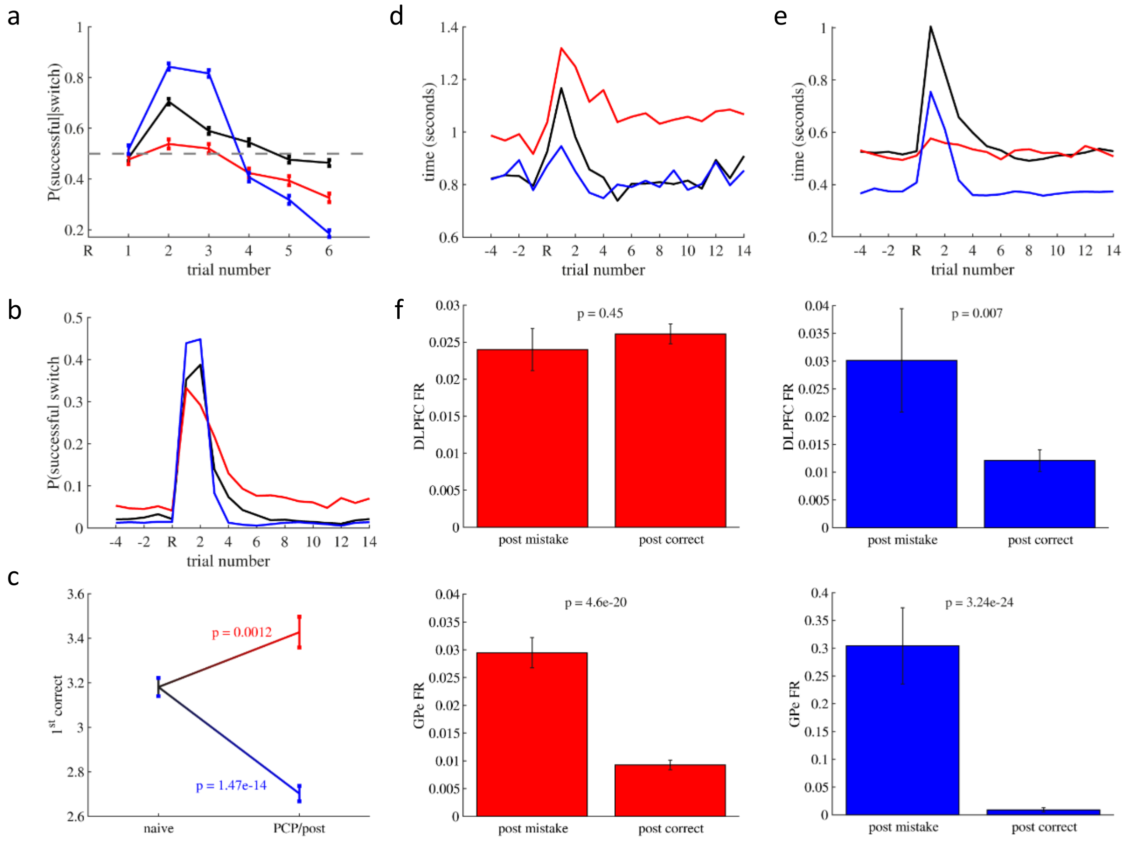


**Extended data figure 7: behavioral changes under PCP administration and post PCP clearance. (a)** Probability of choosing successfully, given that a switch was made in the naïve state (black), under PCP administration (red) and post-PCP (blue). Dashed grey line marks the chance level (P = 0.5). **(b)** Probability of making a successful switch (out of all possible choices). **(c)** number of trials from reversal until choosing the correct stimulus (for the first time), naïve (black), under PCP administration (red) and post-PCP (blue). **(d)** Response time from stimulus choice cue appearance until pressed. **(e)** Response time from reward cue appearance until pressed. **(f)** Mean and SEM of the DLPFC (top) and GPe (bottom) firing rate leading to stimulus choice selection in trial following mistake (post-mistake) and trials following correct (post-correct). Ps represent the p-values corresponding to the two-sample t-test between the mean FR of post-mistake and post-correct trials in each state (PCP and post-PCP). **Left column** – PCP results. **Right column –** post-PCP results.

**Related to main text figure 2.**

### Extended data figures

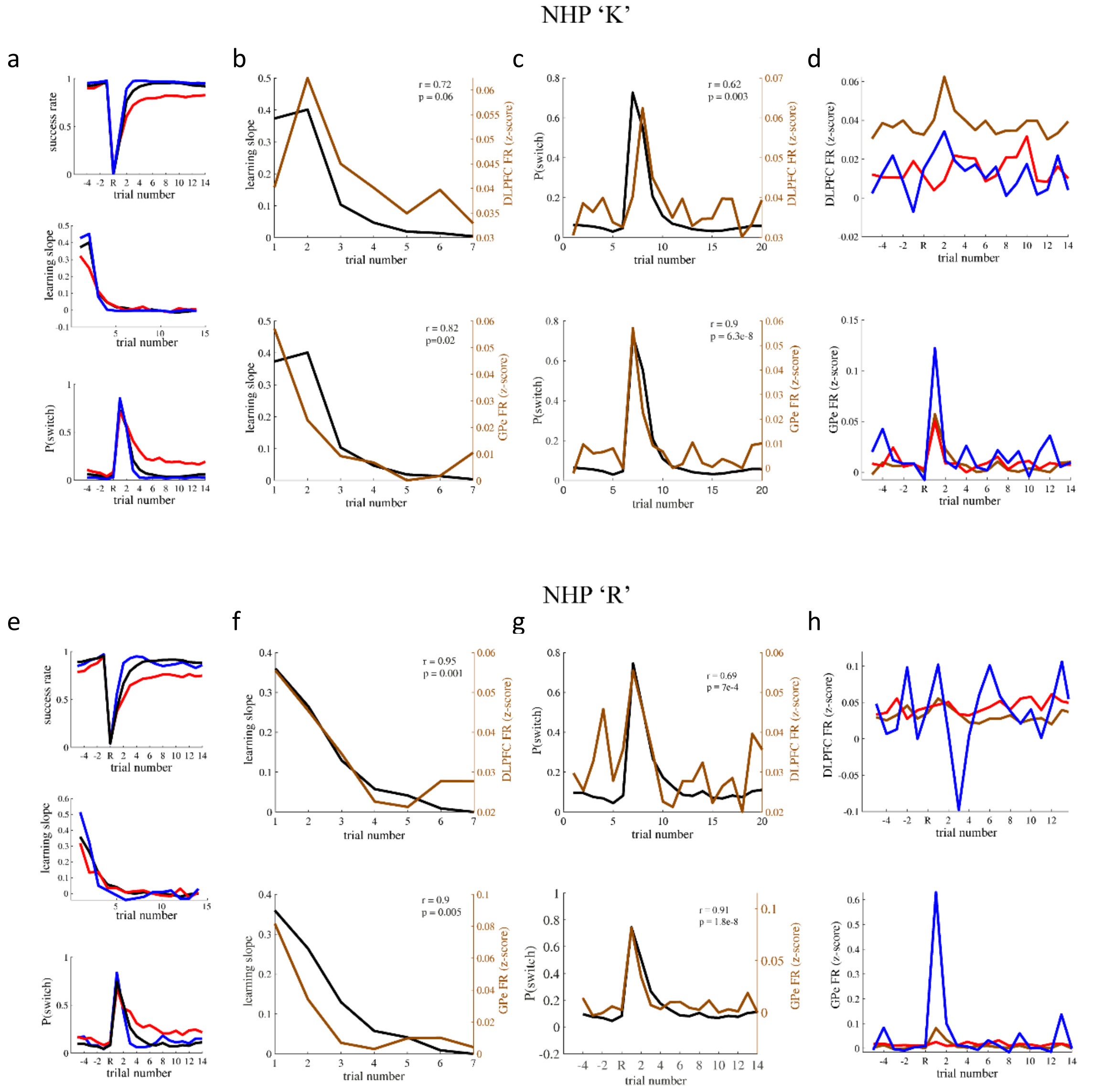


**Figure 8: Both NHPs showed similar neuro-behavioral results across all conditions. (a-d)** NHP K data analysis. **(a)** Behavioral results of all three states: naïve (black), PCP (red), and post-PCP (blue). **Top** – learning curve, **middle** – learning slope and **bottom** – switch probability. **(b)** The learning slope (black) and the DLPFC (top, brown) and GPe (bottom, brown) mean firing rate during the two seconds preceding choice selection. **(c)** switch probability (black) and the DLPFC (top, brown) and GPe (bottom, brown) mean firing rate during the two seconds preceding choice selection. **(d)** Mean DLPFC (top) and GPe (bottom) firing rate during the two seconds preceding choice selection throughout the task of all three conditions: naïve (brown), PCP (red), and post-PCP (blue). **(e-h)** The same as a-d for NHP 'R'.

**Related to main text figure 2.**

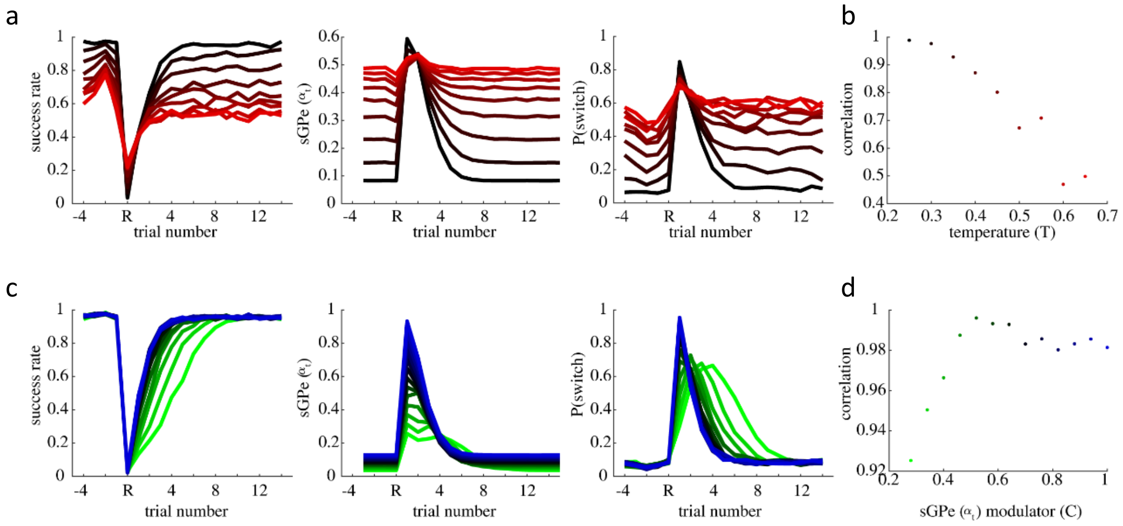


**Extended data figure 9: simulation parameters effects. (a-b)** Simulating the effect of increasing the model's temperature parameter, from black (low temperature) to red (high temperature). **(a)** From left to right – success rate, sGPe value and switch probability during task progression. **(b)** Correlation coefficient value between sGPe value and switch probability. **(c-d)** Simulating the effect of changing sGPe modulator (C) value (i.e., increasing and decreasing the baseline value of sGPe 'activity'). **(a)** from left to right – success rate, sGPe value and switch probability during task progression. **(d)** Correlation coefficient value between sGPe value and switch probability.

**Related to main text figure 3.**

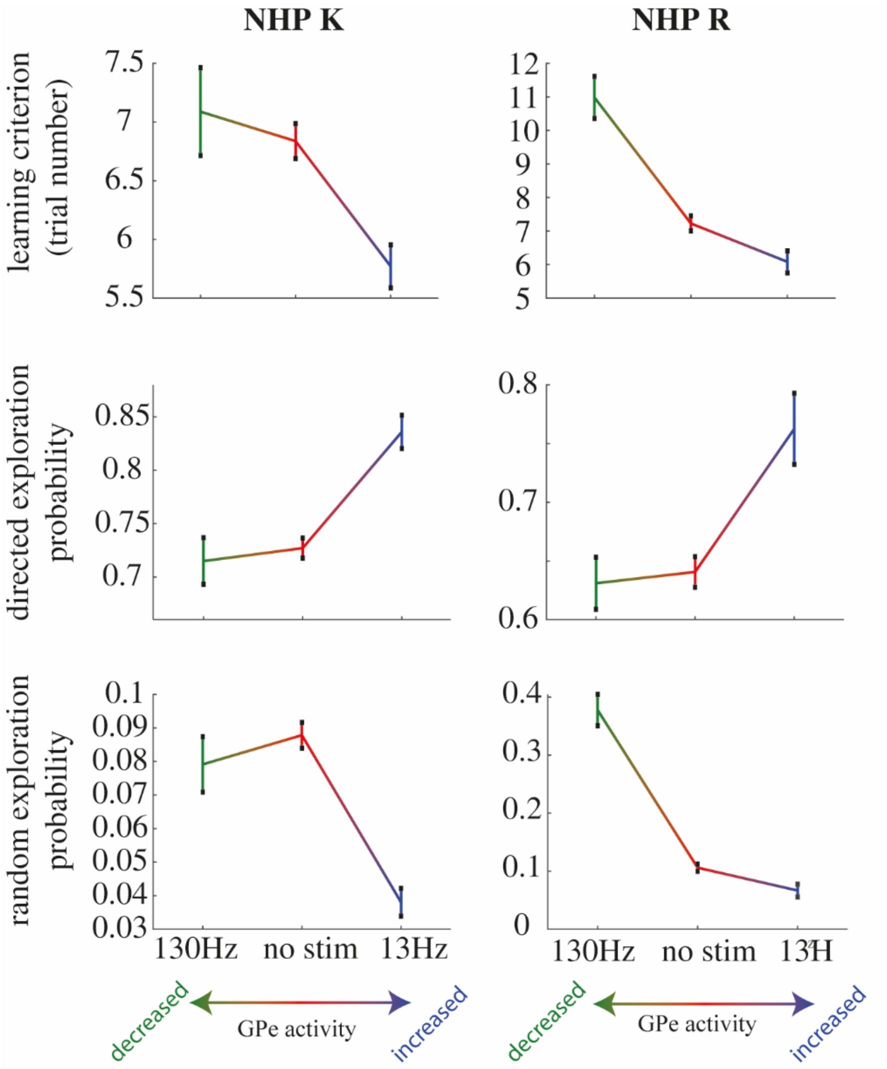


**Extended data figure 10: GPe macro stimulation had similar effect on both NHPs. Left -** NHP K and **Right –** NHP R. **Top -** the NHPs' learning speeds. **Middle -** Probability of initiating directed exploraiton. **Bottom -** Probability of initiating random exploration.

**Related to main text figure 4.**
